# Protein Abundance Prediction Through Machine Learning Methods

**DOI:** 10.1101/2020.09.17.302182

**Authors:** Mauricio Ferreira, Rafaela Ventorim, Eduardo Almeida, Sabrina Silveira, Wendel Silveira

## Abstract

Proteins are responsible for most physiological processes, and their abundance provides crucial information for systems biology research. However, absolute protein quantification, as determined by mass spectrometry, still has limitations in capturing the protein pool. Protein abundance is impacted by translation kinetics, which rely on features of codons. In this study, we evaluated the effect of codon usage bias of genes on protein abundance. Notably, we observed differences regarding codon usage patterns between genes coding for highly abundant proteins and genes coding for less abundant proteins. Analysis of synonymous codon usage and evolutionary selection showed a clear split between the two groups. Our machine learning models predicted protein abundances from codon usage metrics with remarkable accuracy, achieving R^2^ values higher than previously reported in the literature. Upon integration of the predicted protein abundance in enzyme-constrained genome-scale metabolic models, the simulated phenotypes closely matched experimental data, which demonstrates that our predictive models are valuable tools for systems metabolic engineering approaches.

## INTRODUCTION

Proteins are the primary molecules of cellular function. The efficient allocation of the cellular proteome is responsible for controlling metabolic flux and many physiological processes (1). The absolute abundance of each protein is valuable information for genome-scale metabolic reconstructions, as it improves the modelling of metabolic flux and protein allocation (2). Current mass spectrometry technology and quantitative proteomics analysis have allowed the quantification of thousands of proteins in different organisms (3). However, a large portion of proteins are still undetected (4) due to variation in their physicochemical properties, signal intensities, and ionisation efficiencies (5). The high-cost of reagents and equipment is another drawback (6). Absolute quantification has been mostly limited to model species, which hinders systems biology endeavours in non-model species (7). Genome-scale metabolic models (GEMs), which account for protein abundance, have been reconstructed for a limited number of species. A recent review of models of metabolism and macromolecular expression (ME-models) reported only 4 ME-models (8), while 108 stoichiometry-only GEMs (M-model) are available on the BiGG repository (9). Likewise, GEMs with enzymatic constraints using kinetics and omics data (GECKO models) have only been reconstructed for *Saccharomyces cerevisiae* (10, 11) and *Bacillus subtilis* (12). The integration of omics data to GEMs, especially protein abundance, can be useful to improve simulations. For example, the *S. cerevisiae* iMM904 model, which is integrated with proteomic measurements and solved by Linear Bound Flux Balance Analysis (LBFBA), matched more closely experimental fluxomics data than the iMM904 model without proteomics data solved by Parsimonious Flux Balance Analysis (pFBA) (13). This finding highlights the importance of further increasing the number of GEMs integrated with protein abundance.

The abundance of proteins is primarily determined by a combination of factors, such as mRNA abundance, translation efficiency, protein turnover rate, and codon usage bias (CUB) (14). CUB, which is a phenomenon in which certain synonymous codons are employed more frequently than other codons (15), is positively correlated with protein abundance (16). Codons can be optimal or non-optimal, depending on their average decoding time (the time needed for a ribosome to read it) (17), and frequent and rare, depending on how often a certain codon appears in a coding sequence (CDS) (18). Based on this description, codons can be classified as frequent and optimal (FreO), frequent and non-optimal (FreNO), rare and optimal (RareO), and rare and non-optimal (RareNO). The distribution of codons regarding optimality and frequency in a protein-coding sequence is not stochastic, that is, it follows an evolution-selected distribution given their individual contributions to protein biosynthesis (19). For instance, 5’- and 3’-extremities of a CDS have a strong selection against uniformity, in sharp contrast with more central regions. The pattern of codon composition also impacts protein structures, as certain secondary structures, such as coil regions, have an enrichment of RareNO codons that is not detected in other types of structures. The 5’-extremity is also enriched with RareNO codons, with an average decoding rate that is compatible with ramp theory (20). Furthermore, enzymes from central metabolic pathways are highly abundant and present strong codon usage bias (i.e., defined pattern of codon usage). Proteins from stress response pathways, on the other hand, are less abundant and have weaker codon usage bias (i.e., codons are uniformly employed) (17, 21).

As protein abundance is generally conserved across diverse phylogenetic taxa (22), this poses the question of whether absolute protein abundance can be mathematically predicted based on existing data. Based on the association between protein abundance and codon usage bias, it is reasonable to consider that metrics of codon usage bias are a potential source of information that can be applied to predict the abundance of proteins of non-model species or proteins undetected by mass spectrometry.

Machine learning has been a useful tool for systems biology, with applications such as the prediction of enzyme turnover numbers (23), prediction of metabolomics time-series data from proteomics time-series data (24), automated metabolic model ensemble-driven elimination of uncertainty with statistical learning (25), and characterisation and reduction of uncertainty in kinetic models (26). Thus, we hypothesise that machine learning algorithms can predict protein abundances by using codon usage metrics as features.

In this study, we explore the evolutionary constraints that shape the codon usage bias of the *Saccharomyces cerevisiae* genome in the context of protein abundance by comparing highly abundant proteins (HAP) and lowly abundant proteins (LAP). We then apply supervised learning algorithms to a series of codon usage metrics calculated from protein-coding sequences with known protein abundance to predict the abundance of proteins employed by enzyme-constrained genome-scale metabolic models (ecGEMs), including previously undetected proteins. The integration of these predictions in ecGEMs enables phenotype simulations that match those performed with experimentally measured protein abundances.

## MATERIAL AND METHODS

### Data collection

The protein abundance values were obtained from a unified data set of *Saccharomyces cerevisiae* proteome quantifications, which were compiled by Ho et al. (27). This dataset has absolute abundance values that exceed 5,000 proteins; it is composed of absolute and relative measurements from 21 quantitative proteomics analyses. For all analyses, we applied the median absolute abundance values after filtering for GFP autofluorescence. Additionally, we also employed median values for measurements performed using either the minimal medium or the YPD medium.

We retrieved the CDS associated with each protein reported in the proteomics study from Ensembl Fungi (28), using the BioMart tool (29). We obtained other information, such as amino acid sequences, size and molecular weight from UniProt (30). For the tRNA analysis, we recovered the *S. cerevisiae* tRNA gene copy numbers from GtRNAdb (31, 32).

### Characterisation of codon usage

To characterise codon usage bias in the *S. cerevisiae* genome, we assessed how evolutionary constraints affect codon distribution and protein biosynthesis regarding protein abundance. Of the 5,388 proteins with reported absolute abundance values, we chose the top 10% of proteins (538 proteins) with the highest overall median abundances and the top 10% proteins with the lowest overall median abundances. Proteins whose number of molecules per cell was less than 100 and were detected by only one proteomics study were not considered. We analysed the attributes of codons, such as frequency, optimality, and positional dependency, for the CDSs of proteins retrieved from the previously mentioned data set.

First, we analysed the codon positional dependency using the CodG package (19) to assess how different positions in the sequences are under evolutionary selective pressure. We employ a binning scheme, as described by Villada et al. (19), where codons in a sequence are divided in a binning scheme relative to the CDS length. Each CDS is divided into 10 parts, which includes information about codon quantity per tenth part. We generated a matrix of 59 rows, which includes all codons, except start, stop and tryptophan (since it has only one codon), and 10 columns that correspond to the 10 parts. Using this matrix, we tested the codon distribution uniformity by calculating the χ^2^ value of each codon, as described by Hockenberry et al. (33), using the following equation:

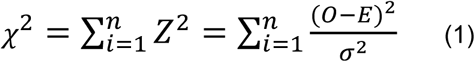

where *O* is the observed count per bin in the group of sequences that are being tested, *E* is the expected value, and *σ* is the standard deviation of the codon counts per bin obtained from 200 simulated groups of sequences, where each amino acid sequence of each CDS in each group (highly or lowly abundant proteins) was conserved but the codons were randomised. The significance of each codon is reported by a p-value of 1.7E-4 after a Bonferroni correction for 59 tests, where the p-value = 0.01, and contrasts them with the χ^2^ distributions, which have *n* - 1 degrees of freedom. For the codon frequency, we calculated the Relative Synonymous Codon Usage (RSCU) (18) using the following equation:

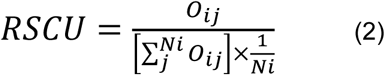

where *O*_*ij*_ is the frequency of the *j-*th codon for the *i-*th amino acid, and *Ni* is the total number of synonymous codons for the *i-*th amino acid. According to Nasrullah et al. (34), codons with RSCU ≥ 1 are considered frequent codons, and codons with RSCU < 1 are considered rare. We also performed principal component analysis (PCA) to identify the distance or relatedness of RSCU values using Orange3 data mining software (35). Next, we calculated the selection on transcript translational efficiency (St) and selection on transcript biosynthetic cost (Sc), as described by Seward & Kelly (36, 37). Subsequently, we performed the Akashi Test (38) to calculate the selection on translation accuracy using the software Seforta (39).

### Feature compilation for machine learning

To evaluate whether machine learning could capture any underlying pattern and predict absolute abundance values using codon usage metrics as features, we compiled a set of features to build predictive machine learning models. We selected codon usage metrics that are calculated individually for each gene. For each CDS, we calculated various codon usage metrics (Table S1) by using EMBOSS (40), CodonW (41), CAIcal (42), coRdon (43), stAIcalc (44), and scripts included in the original manuscripts, such as CodonMuSe (37) and iCUB (45). For codon usage metrics that require highly expressed genes as a reference, we employed the CDSs of the top 10% proteins with the highest overall median abundance.

### Data-set construction

We compiled three separate datasets for training (Figure 1). The first data set uses the median absolute abundances of all 21 quantitative proteomics analyses described by Ho et al. (27) and has a total of 5,388 instances. The second training data set employs the median absolute abundances of experiments that quantified proteins of *S. cerevisiae* yeast growing in minimal media. Eleven quantification experiments were taken into account (abbreviated by Ho et al. (27) as PENG, LAW, LAHT, DGD, THAK, TKA, BRE, DEN, MAZ, CHO, and YOF), which generated 5,187 instances. The third training data set is obtained from experiments in which *S. cerevisiae* is grown in YPD medium (LU, KUL, LEE2, NAG, PIC, WEB, NEW, LEE, DAV, GHA); 5,114 instances are generated. The codon usage metrics listed in Table S1 yielded a total of 91 features, including individual gene codon usage metrics and nucleotide composition numbers (Table S2 and Table S3). After compiling the training data sets, we log-transformed the protein abundance values for the three data sets to favour a normal distribution.

**Figure 1.**
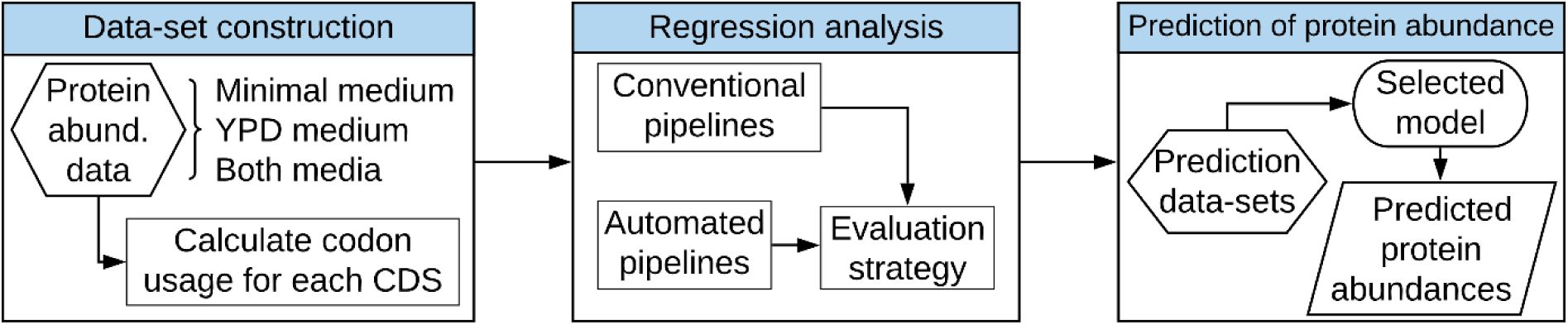
Machine learning applied to codon usage bias for the prediction of protein abundance. We calculated a series of codon usage metrics and nucleotide composition numbers. Three separate data sets were compiled and employed for the training: the first data set contains protein measurements from yeast cultivation in the minimal medium, the second data set consists of protein measurements from yeast cultivation in the YPD medium, and the third data set combines protein measurements for both culture media. The hexagonal boxes denote the input data; the rectangular boxes indicate the intermediate steps; the ellipsoids represent the trained regression models; and the parallelogram denotes the output from these models.

### Regression model training

We explored different machine learning libraries to build a predictive model, namely, Scikit-Learn (46), H2O (47), and XGBoost (48). Since the objective was to predict the absolute abundance of proteins (numerical value), we employed all available algorithms with support for regression problems in the previously mentioned libraries, such as linear models, ensemble models, and neural networks. We also applied automated machine learning pipelines from H2O, TPOT (49), and GAMA (50). The H2O automated machine learning tool trains and cross-validates pre-configured algorithms included in the library to select the best algorithm. TPOT and GAMA utilize genetic algorithms to explore many different possible pipelines using Scikit-Learn algorithms to identify the best pipeline. TPOT also exports the code of its predicted pipeline. We describe the best algorithm for each training data set in the Supplementary Appendix. All the source code was written in Python 3.7 (51).

For manually configured pipelines, we randomly split the data set into two subsets and use 75% for training and 25% for testing. To determine the hyperparameters of each regressor, we applied a randomized approach, where each parameter is sampled from a set of possible parameter values. We selected the values that produced models with the best evaluation metrics when we employed the test data set as input. For the automated ML pipelines, we ran H2O for a maximum runtime of 6 hours with an unlimited number of tested models. We ran TPOT with 1,000 generations and a population size of 250. For GAMA, we employed an asynchronous evolutionary optimization algorithm with a maximum runtime of 3,600 seconds. We trained all automated pipelines with 10-fold cross-validation, where the data set is partitioned into 10 equally sized subsets. One of these subsets is randomly chosen as the test data set; this process is repeated until each subset has been utilized for a test exactly once.

### Model evaluation and selection

After training the models, we evaluated their performance with a separate data set using data that had not been selected for training. We predicted the absolute abundance values of the enzymes integrated in the ecYeast7 ecGEM and compared them to the median overall abundance data from Ho et al. (27). We checked the median absolute deviation (MAD) and the coefficient of determination (R^2^). The MAD is defined as the median value of all absolute differences between the predicted values and the real values. We chose this metric since the absolute abundance of proteins employed for training are expressed as a median and it is robust to outliers. This metric can be calculated according to the following equation:

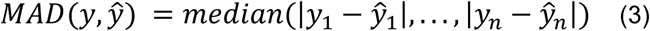

where ŷ_*i*_ is the predicted value of the _*i*_-th sample and *y*_*i*_ is the known protein abundance value. When evaluating the predictive models, we searched for models with the smallest possible MAD. R^2^ is a measure that represents the variance explained by independent values in the regression model; it is calculated as

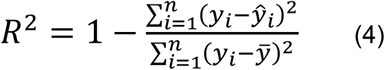

where *y*_*i*_ is the known protein abundance value, ŷ_*i*_ is the predicted value, and 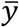 is the mean of the observed data. We searched for models with an R^2^ equal to or greater than 0.75. Models with a high MAD and low R^2^ were re-trained with adjusted hyperparameters until they converged. We selected the model for each data set with the highest R^2^ and lowest MAD for the next step.

### Integration of predicted abundances in the enzyme-constrained metabolic model ecYeast8

After selecting the best predictive models, we decided to predict the abundance of all enzymes employed by the ecYeast8 ecGEM, which has 968 enzymes integrated with enzyme turnover values (*kcat*) but no integration of protein abundance values. Codon usage metrics were calculated for the CDS of all 968 enzymes. We compiled three prediction data sets in accordance to the medium culture utilized for protein quantifications: minimal medium, YPD medium, and a combination of these two media. The difference between each data set is the reference highly expressed genes applied to calculate the codon usage metrics, as described in the feature compilation section.

To validate the predictions obtained by machine learning, we integrated the predicted protein abundances in the ecYeast8 model. We applied the GECKO Toolbox (10) to set the protein abundances as the upper bound for reactions that use enzymes. Considering that the predicted abundance values are expressed as the number of molecules per cell and the ecGEM requires protein abundance values in millimoles per gram of cell dry weight (mmol/gDW), the values were converted to mmol/gDW by assuming a total cellular protein mass of 0.448 gram of protein per gram of cell dry weight (g/gDW), a cellular density of 1.5 × 10^7^ cells per litre (cells/L), and a total biomass of 3 grams of cells dry weight per litre (gDW/L). These values were obtained from (10), (52), and (53), respectively. We re-fitted the parameters *f, GAM* and *sigma* using the batch model before constraining the enzymes. After integrating the protein abundances, the upper bound of the enzyme-constrained reactions was flexibilized to optimize growth at a dilution rate of 0.1 (h^−1^) by setting the carbon source as glucose with an uptake rate of 1.1 mmol/gDWh^−1^. To check whether the unit conversion step was performed correctly, we also re-ran the enzyme abundance integration step with ecYeast7 using the quantitative proteomics data from Lahtvee et al. (54).

### Model growth simulations

We attempted to replicate the results obtained by Sánchez et al. (10) and compared the simulations of the Yeast7 model with the ecYeast7 model using experimental data obtained from (55). We simulated a chemostat with a dilution rate fixed at 0.1 h^−1^. We removed any constraints for substrate uptake and limited unmeasured enzyme mass by 0.448 g/gDW, set the *f* value to 0.4421 g/g, and used a *σ* value of 0.5. For the optimization, we minimized the glucose uptake rate and fixed the glucose uptake rate to the optimal value with a 0.1% flexibility, and minimized enzyme usage. Regarding the ecYeast8 model integrated with ML-predicted abundances, we simulated a chemostat with the same previously mentioned conditions, except that unmeasured enzyme mass was not limited, as the protein pool pseudo-reaction was not included in the models. We compared the results to experimental fluxes measured by ^13^C-MFA obtained from (55). For all simulations, we checked the metabolic fluxes regarding consumption of glucose and O_2_ and the production of CO_2_. We have also compared the models by executing flux variability analysis (FVA). All model configuration and problem setups were performed with the RAVEN Toolbox 2 (56) in MATLAB 2017a (The MathWorks Inc., Natick, Massachusetts). Problems were solved using the Gurobi Optimizer version 8.11 (57).

## RESULTS

### Codon usage is markedly different between the coding sequences of highly abundant proteins and those of lowly abundant proteins

We evaluated how evolutionary constraints shape the codon usage bias of the *S. cerevisiae* genome by comparing the CDSs of HAP and LAP. We observed a noticeable contrast between the CDSs of HAP and those of LAP. Regarding the codon frequency, in the principal component analysis of RSCU values, two distinct groups of CDSs were observed. The first group is composed of mostly HAP, whilst the second group consists of mostly LAP (Figure S1). A heatmap of the 20 most and least abundant proteins showed that codons of HAP CDSs are more enriched with frequent codons and that rare codons are totally or almost totally depleted (RSCU ≈ 0). Meanwhile, CDSs of LAP have a weaker bias in codon usage, that is, there is no preference for certain synonymous codons (Figure 2).

**Figure 2.**
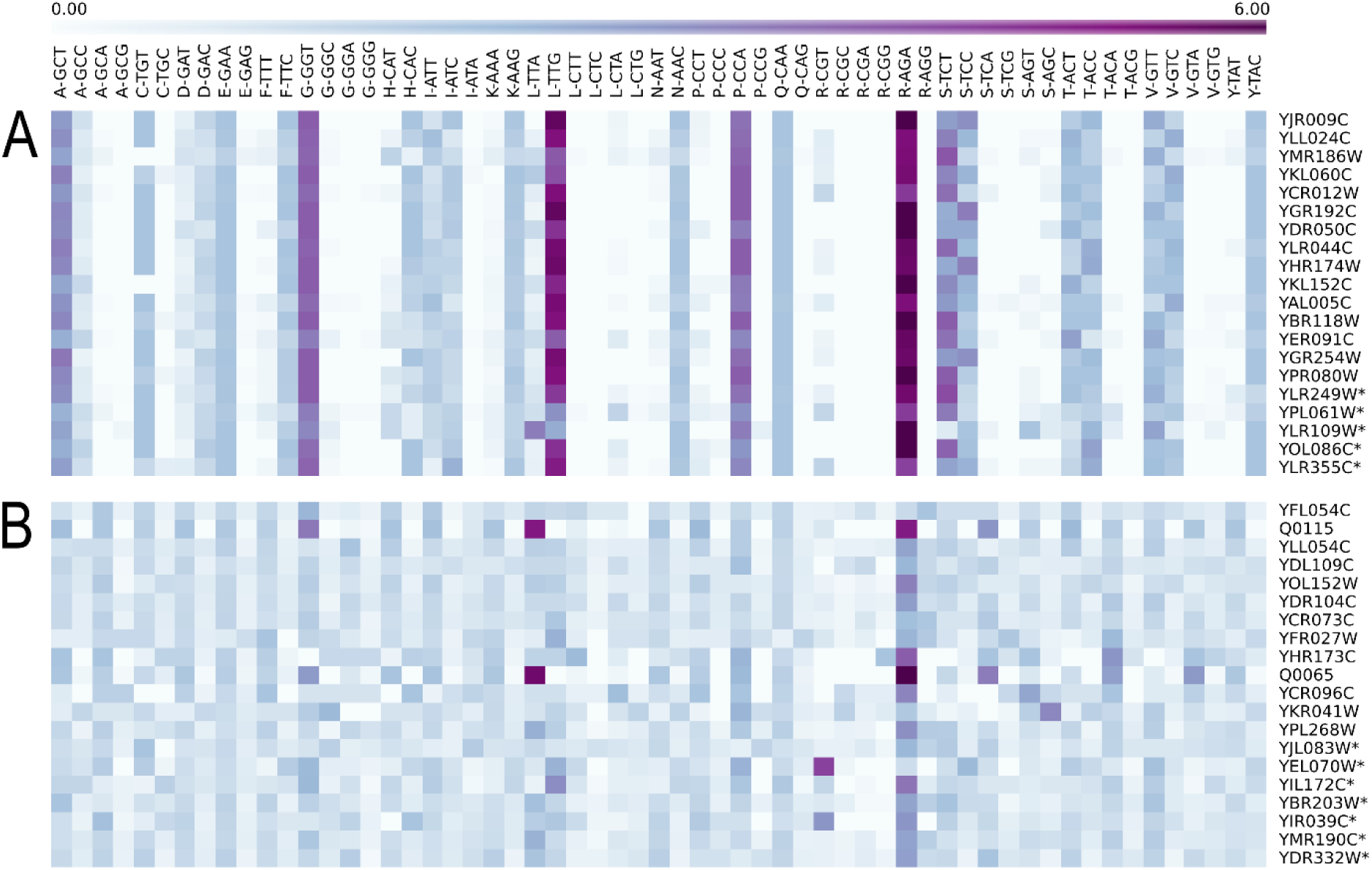
Heatmap of relative synonymous codon usage that illustrates the difference in codon usage in regard to frequency between the 20 most abundant and 20 least abundant proteins of *S. cerevisiae*. Columns denote the 59 codons with one or more synonymous codons. (A) Rows indicate CDSs of the 20 most abundant proteins. (B) Rows represent CDSs of the 20 least abundant proteins. CDSs of the HAP, in contrast to the CDSs of LAP, have many enriched (high RSCU) or depleted (low RSCU) codons. Sequences marked with an asterisk denote the presence of a signal peptide at the 5’ extremity as detected by SignalP 5 (58).

For the top 10% proteins with the highest abundances (n = 538), we observed selection against uniformity at the 5’ end in the CDSs, as indicated by the enriched codons in the first bin. Otherwise, CDS encoding for LAPs (top 10% lowest, n = 538) presented a higher uniformity of codon distribution (Figure 3 and Figure S2).

**Figure 3.**
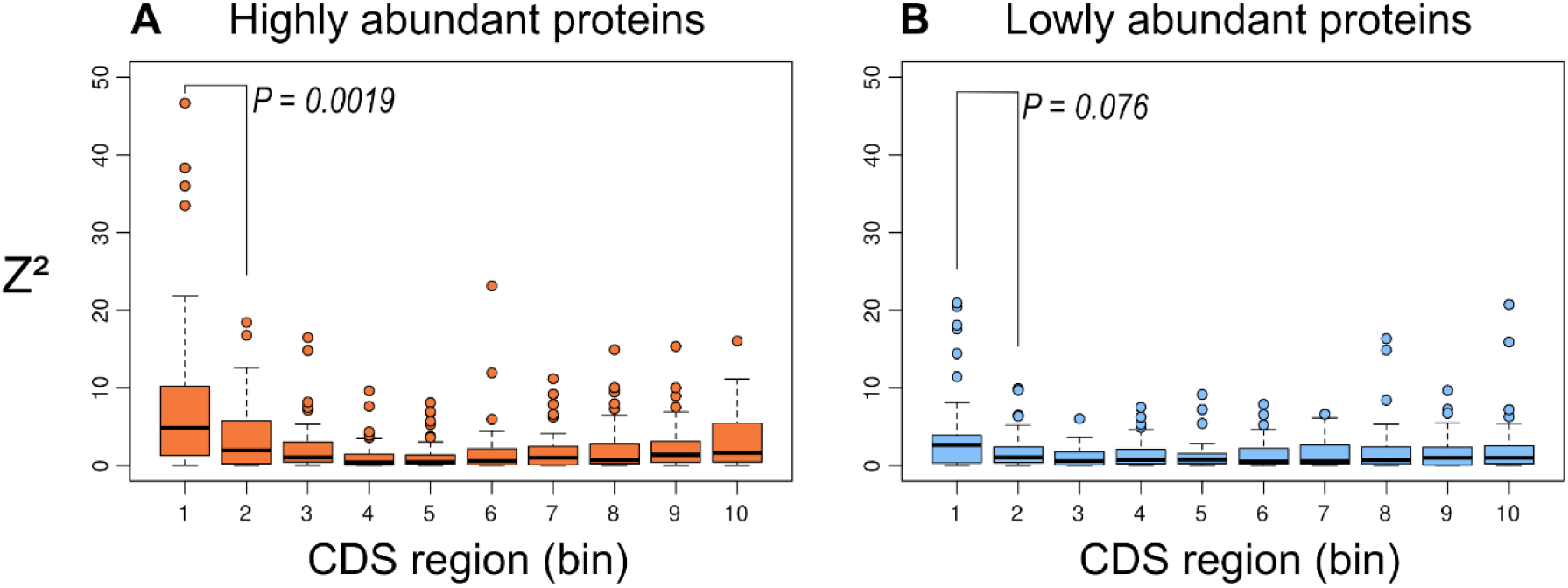
Evolutionary selection for position-dependent codon usage bias determined by the chi-squared test for the matrix constructed with the CodG package. Each CDS was equally divided into 10 bins to evaluate how each position contributes to overall codon usage. (A) Deviation from uniformity in the CDSs of HAPs shows bias towards 5’ end. (B) CDSs of lowly abundant proteins show higher uniformity.

Since the values of translation efficiency (St) and selection for biosynthetic costs (Sc) represent how strongly natural selection acts on the translational efficiency and biosynthetic cost of codons, respectively, we calculated them for both HAP and LAP. Importantly, Sc values suggest that HAP CDSs undergo selection pressures to reduce the biosynthetic cost (Sc < 0) and maximize the translation efficiency (St > 0) (Figure 4).

**Figure 4.**
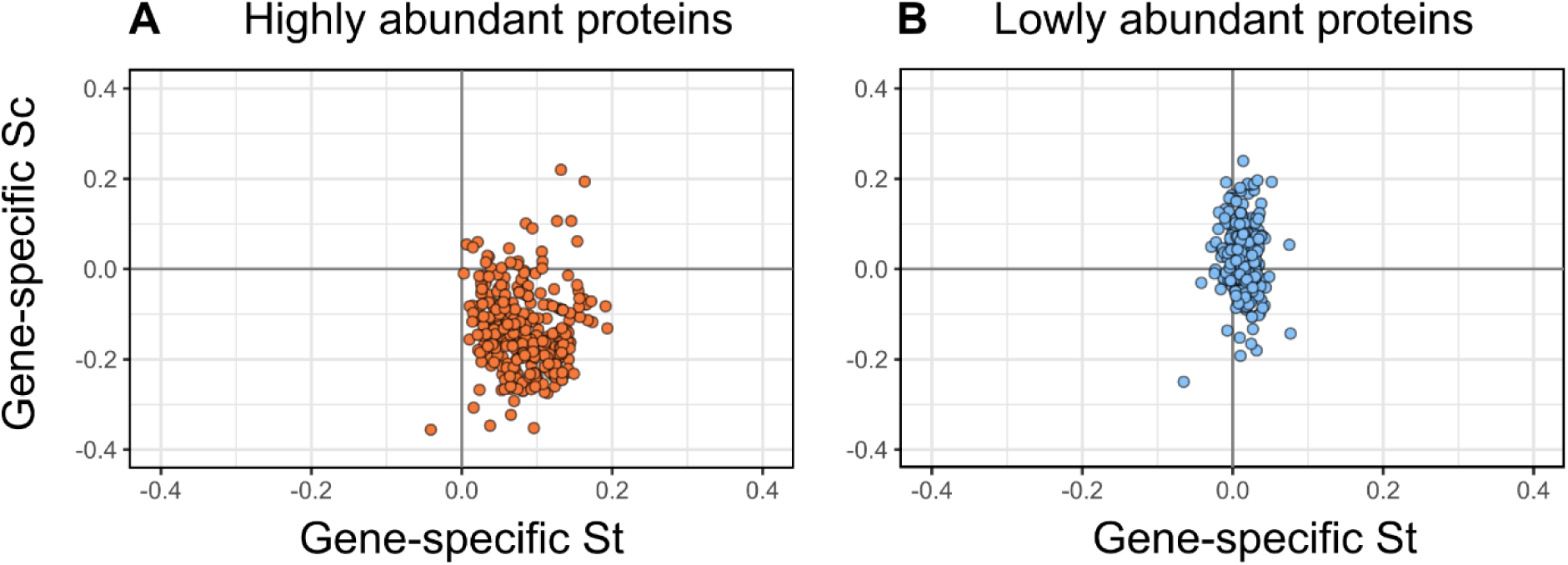
Codon evolutionary selection for translation efficiency (St) and biosynthetic costs (Sc). Higher St values indicate that the gene is strongly selected for translation efficiency, whilst lower Sc values indicate that the gene is strongly selected to reduce the biosynthetic cost of codons. (A) Most Sc values of the CDSs of HAPs are lower than zero, whereas most St values of the CDSs of HAPs are higher than zero, which suggests that they are selected for decreased biosynthetic costs and increased translation efficiency, respectively. (B) CDSs of LAPs did not show any tendency towards any direction.

On the other hand, the LAP sequences did not show any tendency, that is, mean St ≈ 0 and mean Sc ≈ 0. Interestingly, the Akashi test revealed that both groups are subjected to the same selection for translation accuracy (mean odds ratio ≈ 1) (Figure 5). A combination of these results indicate that the codon usage bias is different between the CDSs of LAP and those of HAP.

**Figure 5.**
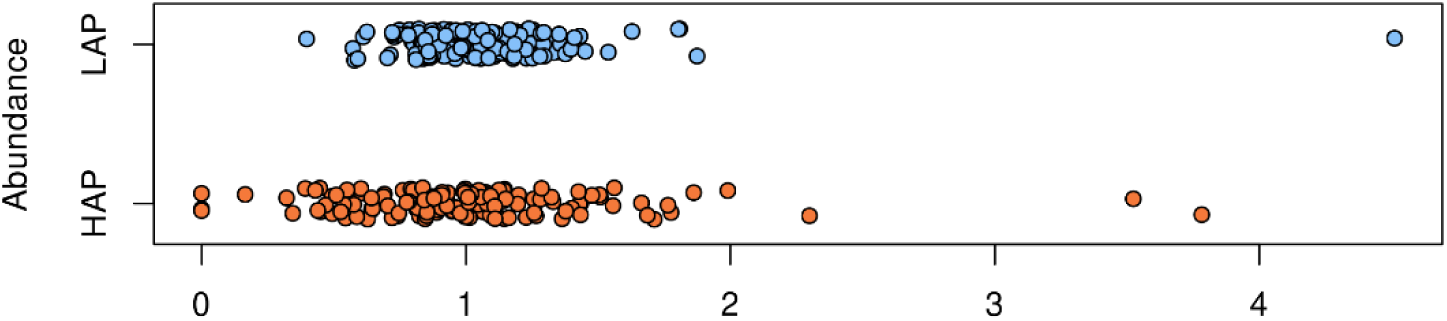
Codon evolutionary selection for translation accuracy. Each dot represents a CDS of HAP or LAP. The CDS of both highly and lowly abundant proteins are centred around an odds ratio of 1, which suggests that both groups are subjected to the same selection for translation accuracy. The odds ratio represents the association between a set of optimal codons and evolutionary constrained sites.

### Machine learning models can predict protein abundance

Once the three data sets were compiled, we separately trained predictive models for each data set. We tested each algorithm with support for the regression problems from three different libraries (Scikit-Learn, H2O, and XGBoost) and tested the automated machine learning pipelines (TPOT, H2O, GAMA). By validating with an independent data set, which contained 456 enzymes with measured protein abundance integrated in the ecYeast7 model, we evaluated the MAD and R^2^ for all tested algorithms of the three libraries. The results for the five best conventional algorithms are shown in Table 1, and those for all tested algorithms are shown in Table S4.

**Table 1.**
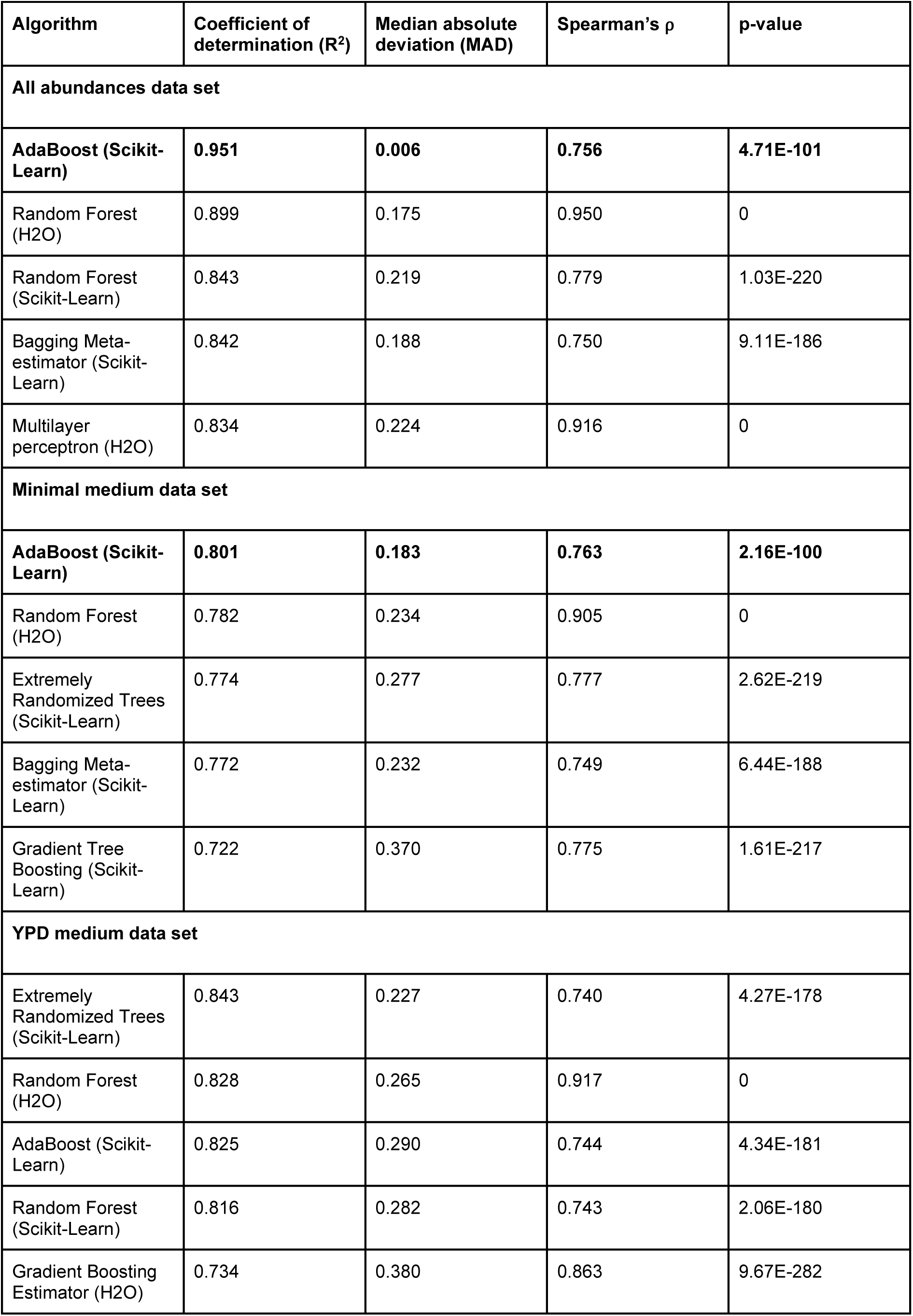
Regression evaluation metrics for the five best trained algorithms for each data set. Each algorithm was trained and evaluated by hold-out validation (75% training, 25% validation). In addition, we employed an independent data set for testing. Spearman’s ρ and its associated p-value assesses the correlation between the predicted values and the median values obtained by Ho et al. (27).

Regarding the automated machine learning tools, the best predicted pipeline for the all media data set was a stacked ensemble predicted by TPOT. For the minimal medium data set, the best predicted pipeline was also a stacked ensemble predicted by H2O; for the YPD medium data set, it was a gradient boosting machine from the XGBoost library predicted by H2O. We observed that the model performances for the automated pipelines were like those for conventional implemented pipelines (refer to Table 2).

**Table 2.**
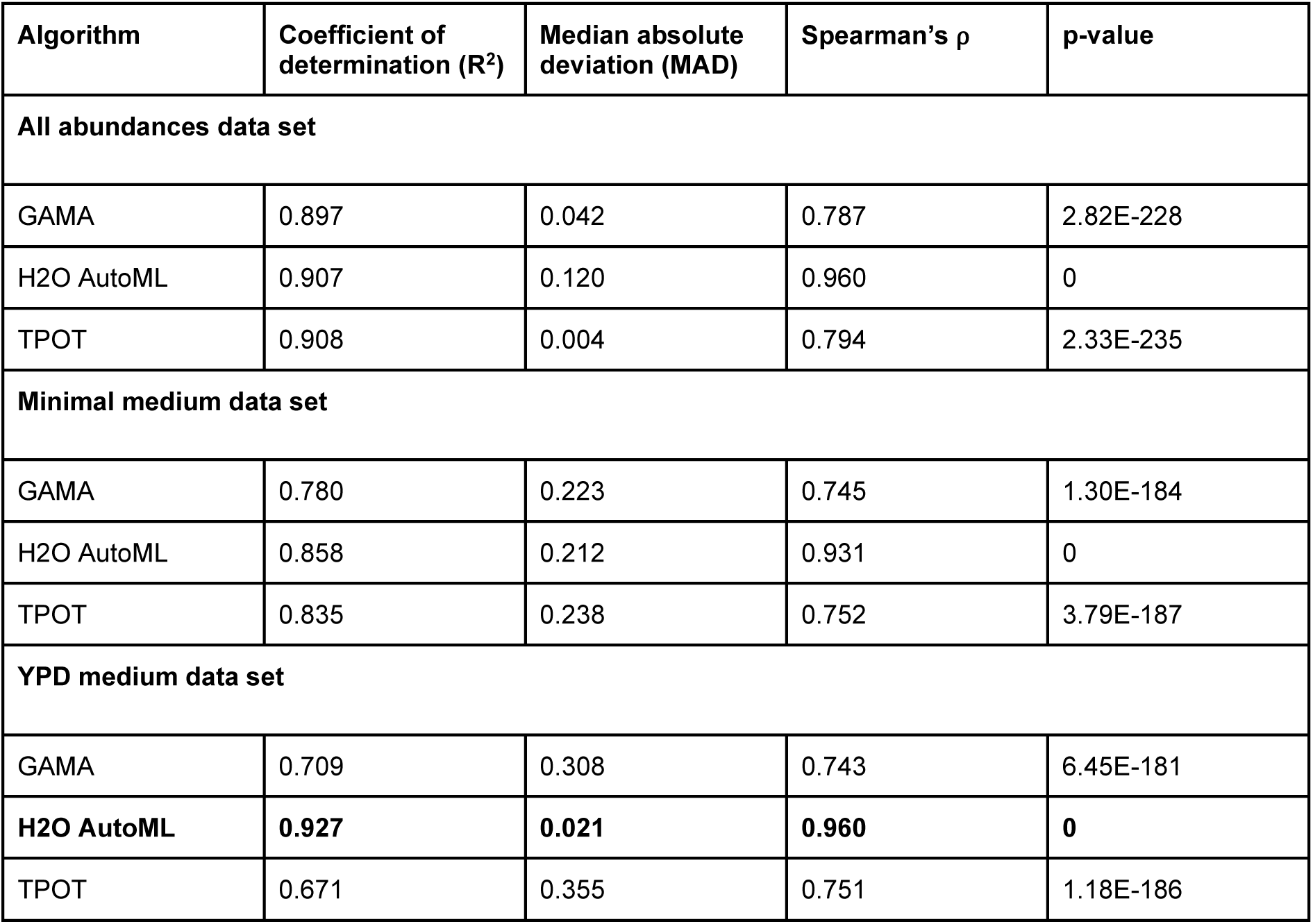
Regression evaluation metrics for automated machine learning pipelines for each data set. Each algorithm was trained and evaluated by 10-fold cross-validation using an independent data set for testing. Spearman’s ρ and its associated p-value assesses the correlation between the predicted values and the median values obtained by Ho et al. (27).

| Algorithm | Coefficient of determination ($R^2$) | Median absolute deviation (MAD) | Spearman's $\rho$ | p-value |
| --- | --- | --- | --- | --- |
| **All abundances data set** |  |  |  |  |
| GAMA | 0.897 | 0.042 | 0.787 | 2.82E-228 |
| H2O AutoML | 0.907 | 0.120 | 0.960 | 0 |
| TPOT | 0.908 | 0.004 | 0.794 | 2.33E-235 |
| **Minimal medium data set** |  |  |  |  |
| GAMA | 0.780 | 0.223 | 0.745 | 1.30E-184 |
| H2O AutoML | 0.858 | 0.212 | 0.931 | 0 |
| TPOT | 0.835 | 0.238 | 0.752 | 3.79E-187 |
| **YPD medium data set** |  |  |  |  |
| GAMA | 0.709 | 0.308 | 0.743 | 6.45E-181 |
| **H2O AutoML** | **0.927** | **0.021** | **0.960** | **0** |
| TPOT | 0.671 | 0.355 | 0.751 | 1.18E-186 |

We determined that the AdaBoost estimator from Scikit-Learn, which was implemented with the TPOT-predicted stacked ensemble as a base estimator, was the best predictive model for the data set of all protein abundances and the data set of the minimal medium abundances; it achieves R^2^ values of 0.951 and 0.801, respectively (refer to Supplementary Appendix for details). For the YPD medium data set, the gradient boosted tree from the XGBoost library, which was predicted and optimized by the H2O automated tool, was the best model with an R^2^ of 0.927 and the lowest MAD for the data set.

### Integration of predictions into enzyme-constrained GEMs

Since we were able to predict the absolute abundance of all 968 enzymes with reasonable accuracy, we decided to incorporate them into the genome-scale metabolic model of *S. cerevisiae*, ecYeast8. We were interested in demonstrating that our predictive model could be useful for reconstructing enzyme-constrained GEMs. For this purpose, we converted the unit from n° of protein molecules/cell (absolute abundance) to mmol/g_DW_, as required for simulating the metabolism (refer to Material and Methods). Using the GECKO toolbox described by Sánchez et al. (10), we obtained three different models—ecYeast8-MIN, ecYeast8-YPD and ecYeast8-ALL—using the minimal medium prediction data set, YPD medium prediction data set, and predicted abundances for all the 21 protein measurements in the prediction data set, respectively. Note that the protein pool pseudo-reaction was not included in the models as we incorporated the abundance for all enzymes.

### Metabolic fluxes simulated with ML-predicted enzyme abundances are similar to experimental data

The reproduction of the results using the ecYeast7 model and quantitative proteomics data (54) showed that our unit conversion step was correctly performed, as it closely approached the original predictions (Figure S3). After including the predicted protein abundances in ecYeast8, we performed a chemostat simulation to compare the performance of the three models to the conventional GEM Yeast8 and the experimental fluxes quantified by ^13^C-MFA. When grown aerobically at a dilution rate of 0.1 h^−1^ and limited by glucose concentrations, all models predicted similar fluxes to the experimental values and the conventional model (Figure 6A). Between the three models, ecYeast8-ALL displayed the best predicted values. The predicted fluxes of both ecYeast8-MIN and ecYeast8-YPD were lower than the fluxes predicted by ecYeast8-ALL. We further compared the models by a flux variability analysis. The three ecGEMs had significant reductions in flux variability for most reactions when compared to Yeast8 (Figure 6B).

**Figure 6.**
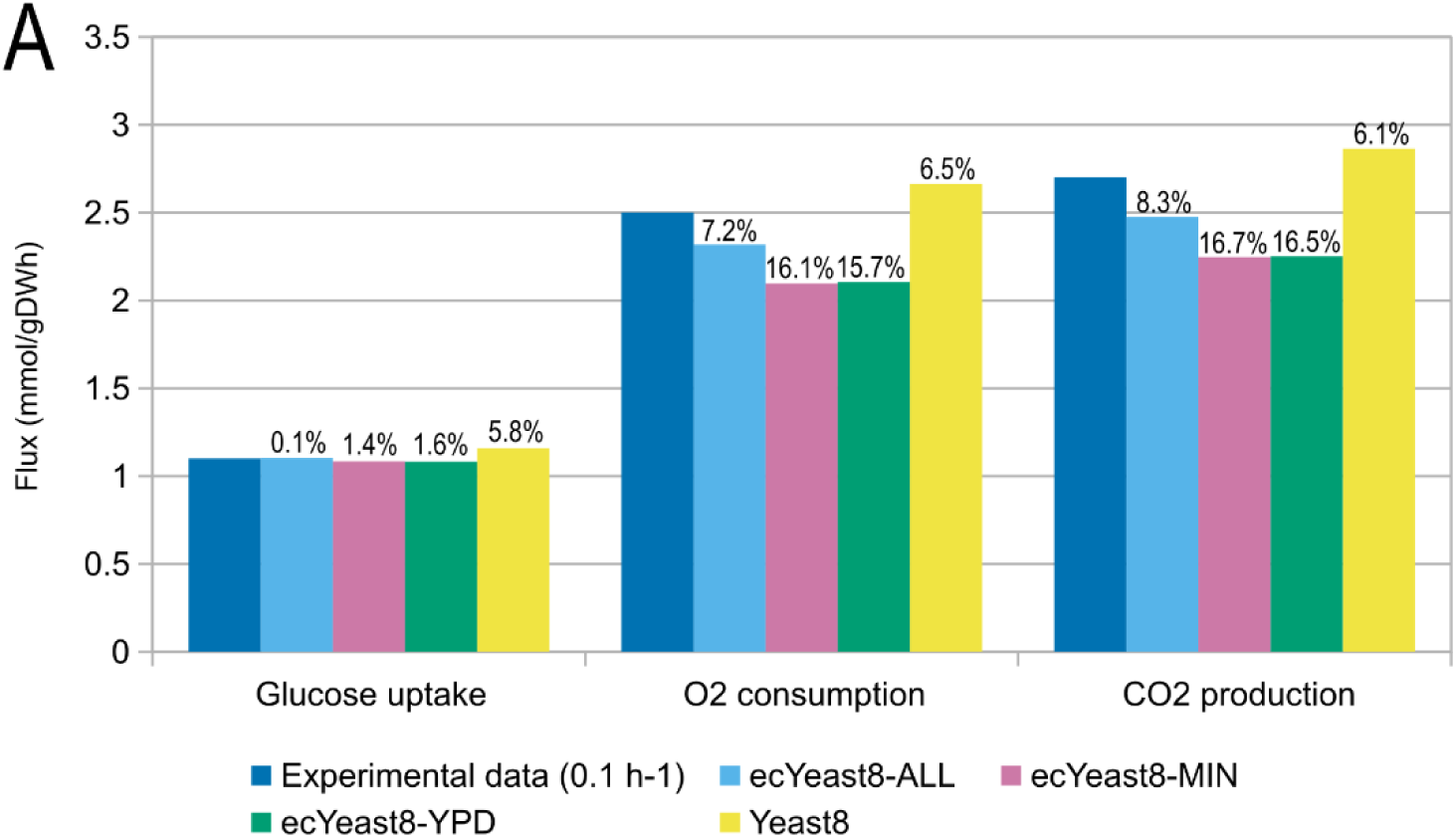
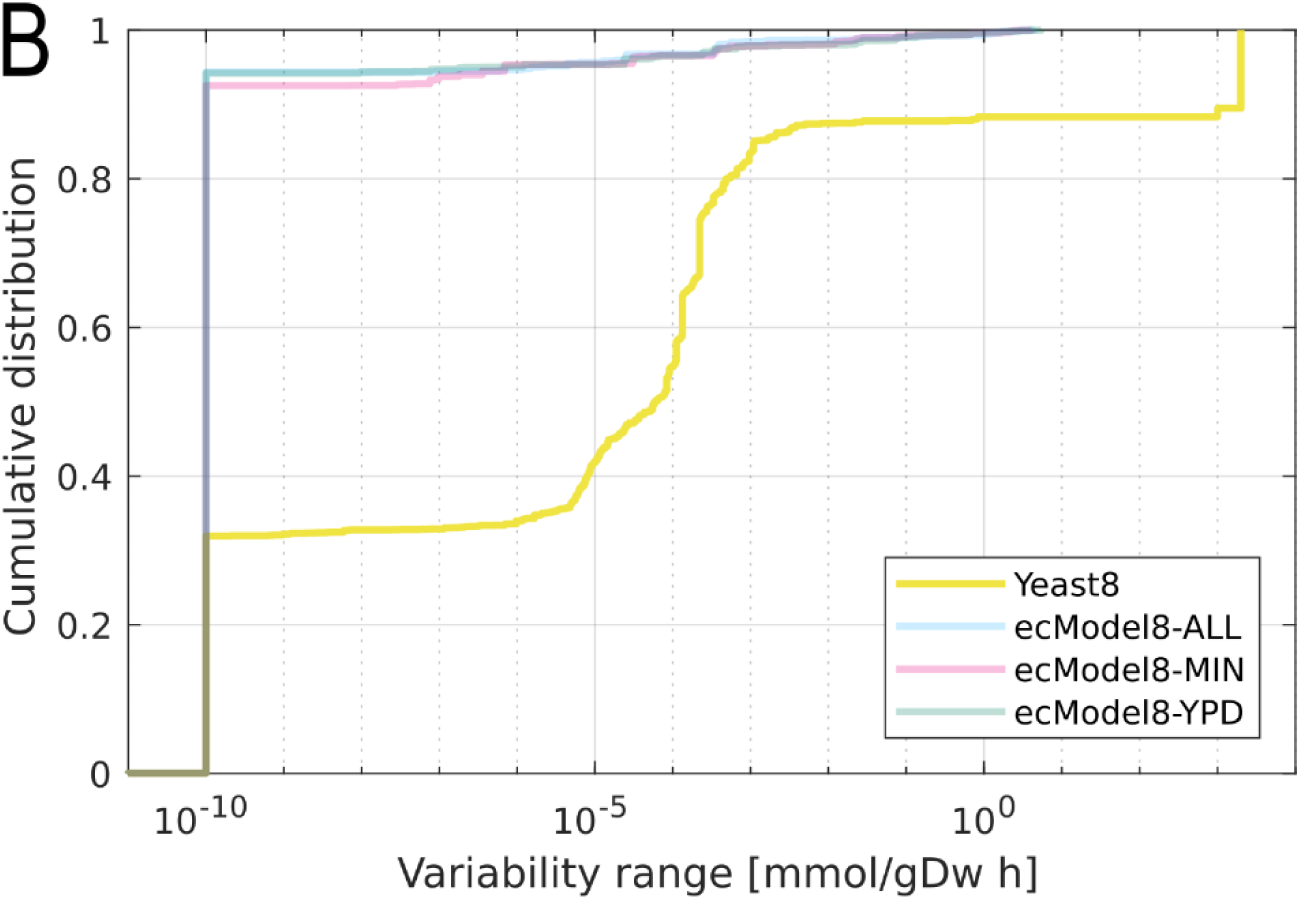
Comparisons between the three ecYeast8 integrated with ML-predicted protein abundances and the traditional GEM Yeast 8. A) Predictions of metabolic flux obtained by the ecYeast8 model integrated with ML-predicted protein abundances and the Yeast8 model (without enzyme constraints), compared to the experimental data. Percentage values represent the relative error when compared to experimental values. B) Flux variability cumulative distribution for Yeast8 and ecYeast8 integrated with ML-predicted protein abundances.

## DISCUSSION

In this study, we explored the evolutionary constraints that affect codon usage bias of CDSs of highly and lowly abundant proteins. The difference between both groups of proteins boosted us to evaluate whether this pattern could be recognized by machine learning algorithms and employed to predict protein abundances. Machine learning algorithms remarkably predicted protein abundances from codon usage metrics. Predicted abundances of proteins were integrated in enzyme-constrained genome-scale metabolic models and successfully applied to simulate metabolic fluxes.

Although synonymous codons can code for the same amino acid, comparative analysis of protein-coding sequences has revealed a “preference” of certain codons. This bias in codon frequency is related to the concentration of tRNA molecules with complementary anticodons and their gene copy numbers (citation). We show that RSCU values separate proteins into two groups: HAP and LAP, which shows that codon usage bias varies according to protein abundance levels. The CDSs of HAP have the highest or lowest RSCU values, which underscores a strong codon bias. Otherwise, the CDSs of LAP have no discernible bias. Consistent with our results, Novoa et al. (59) observed that for a large number of amino acids, codon usage completely changes depending on the expression level.

The stochastic nature of cognate tRNA recognition in the ribosome A-site enables evolutionary forces to act by selecting codons that more closely match the abundance of tRNAs in the cell (60). This selection is likely associated with the differences observed in both translation initiation rates and translation elongation rates (20, 61). The enrichment of certain codons positions of a gene is also determined by natural selection (19), such as the presence of clusters of RareNO codons at the 5’ extremity. We tested this finding by quantifying the position-dependent codon usage bias using a binning scheme as detailed by Villada et al (19). We observed that the CDSs of HAP follows the results obtained from the genome-scale analysis, which show a deviation from uniformity most markedly in the first bin, which was not observed in the CDSs of LAP. This finding agrees with the ramp theory, which poses that a “bottleneck” at the beginning of a CDS is necessary to slow ribosomes to prevent jams and collisions (14, 20).

There are selective pressures regarding codon usage that act on the resource allocation for protein biosynthesis (36, 37), translation efficiency (62), and translation accuracy (38). The resource allocation for protein biosynthesis governs the synthesis of RNA molecules, and polypeptides, as well as other molecules involved in the translational process. One way to reduce the overall biosynthetic cost of a protein without altering its amino acid sequence is to reduce the expenditure on RNA synthesis or increase the translation efficiency of existing RNA molecules (37). Genome-scale analysis of 1,320 bacterial genomes performed by Seward & Kelly (37) revealed that genes subjected to strong selection for reducing biosynthetic cost are also subjected to strong selection to increase the translation efficiency.

In this study, we performed translation efficiency and translation accuracy analyses on CDSs of HAP and LAP and observed that the results rely on protein abundance, which was also observed by Seward & Kelly (37). HAPs experience the strongest selection for reducing the biosynthetic cost and increasing the translation efficiency. By increasing the translation efficiency, more proteins could be synthesized with less mRNA. As observed by Ho et al. (27), the function of HAPs seems to be related to processes such as ribosome biogenesis, protein biosynthesis and cell morphogenesis. The fact that these proteins undergo evolutionary selection for increased translation efficiency and decreased biosynthetic cost is consistent with their high demand by the cell at most physiological conditions to maintain the metabolic activity. On the other hand, LAPs do not seem to be affected by the selection of these parameters. These proteins are associated with processes such as DNA replication, DNA repair, RNA processing and cell-cycle regulation, which are necessary in small amounts and therefore might not require resource allocation optimization through natural selection. Therefore, the difference observed between HAP and LAP in terms of CUB and its relation to the biosynthetic cost reinforces our hypothesis that codon metrics can be useful for predicting protein abundance.

Translation accuracy is also important for reducing the biosynthetic resource cost of a protein and improving translation efficiency, as inaccurate translation elongation increases the time needed to sequester a ribosome, which causes a diminished availability of ribosomes and protein misfolding (63). By applying the Akashi test, we observed that CDSs of both HAP and LAP are subjected to the same strength of selection. This finding was also experimentally noted by Yannai et al. (63) in *Escherichia coli* genes.

As we observed that codon usage bias varies depending on the protein abundance, we were interested in evaluating whether machine learning could capture any underlying pattern and predict absolute abundance values using codon usage metrics as features. We were also interested in addressing whether proteome quantifications with different yeast culture media could impact these predictions. We trained regression algorithms with a series of codon usage metrics for three different data sets and predicted the abundance of proteins that were previously unincorporated in the ecYeast8 model. The data sets differed regarding the type of medium culture employed in each proteomics experiment: we applied the median of the protein measurements obtained for the minimal medium, YPD medium, and combination of both media. After evaluating the regression metrics, we selected the AdaBoost estimator for the data sets of the minimal medium and combination of the minimal and YPD media. The AdaBoost algorithm employs a combination of “weak learners” for the training and predictions; it supports the integration of many other machine learning algorithms as weak learners to improve its performance (64). The TPOT run exported a stacked ensemble of several algorithms; thus, we decided to integrate the predicted pipeline into AdaBoost. We discovered that this approach outperformed all other algorithms and achieved higher R^2^ scores and lower MAD values. For the YPD medium data set, however, this approach was bested by another algorithm. The selected algorithm for this medium was the gradient boosted tree from the XGBoost library, which was predicted and optimized by the H2O automated tool.

There have been many attempts to predict protein abundance values, usually for imputation of abundance values of missing proteins (65–68). Mehdi et al. (68) utilized a Bayesian network that depends primarily on protein abundance and mRNA properties, such as interaction with proteins, expression, and folding energy. Even though these studies obtained satisfactory correlation values, many of their features rely on experimental data. The advantage of our approach is that our features depend entirely on intrinsic information contained in gene sequences (i.e., codon usage metrics), which can all be determined *in silico* and still yield reasonable correlation values. An earlier effort by Nie et al. proposed a zero-inflated Poisson-based model that employed microarray data to predict protein abundance of *Desulfovibrio vulgaris*. Torres-García et al. (66) and Li et al. (67) improved on the previous work by predicting the abundance of *D. vulgaris* proteins using gradient boosted trees (GBT) and neural networks, respectively. However, the R^2^ for both algorithms was lower than that obtained by our study (refer to Tables 1-2). For the GBT algorithm, the R^2^ varied between 0.393 and 0.582, while for the neural networks, it ranged from 0.47 to 0.68. Considering the regression metrics in these comparisons, our proposed prediction models present significant improvement over previous models.

The proposed regression models predicted protein abundances with remarkable accuracy. Thus, we evaluated whether these predictions could be integrated into GEM to simulate the metabolic phenotypes. Comparing the simulated values to the experimental values would be a way to demonstrate the application of our method for constraint-based metabolic modelling. We compared the models based on protein measurements obtained from a minimal (ecYeast8-MIN) medium, rich medium (ecYeast8-YPD) and combination of both media (ecYeast8-ALL). We did not observe differences between ecYeast8-MIN and ecYeast8-YPD. Referring to the results of quantitative proteomics studies, Ho et al. (27) showed that the same medium did not cluster after being subjected to hierarchical clustering or k-means clustering. This finding could explain the similarity between ecYeast8-MIN and ecYeast8-YPD. Notably, the ecYeast8-ALL outperformed the models ecYeast8-MIN and ecYeast8-YPD since its simulations were closer to the experimental values. Consistent with this result, the results obtained using ecYeast8-ALL were similar to the results obtained by Sánchez et al. (10), which were based on experimental protein measurements. Note that the cumulative distribution of the flux variability in the three ecGEMs showed a variability range lower than that for Yeast8, which highlights their capacity to predict metabolic fluxes mostly by reactions constrained by enzymes. According to our results, Sánchez et al. (10) also showed that the inclusion of enzymatic constraints reduced the flux variability of simulations for the ecYeast7 model when compared to Yeast7. The same result was reported for ecYeast8 (11) and the ecGEM of *B. subtilis* ec_iYO844 (12). Importantly, our predictions also maintained a physiologically relevant solution.

Considering that species from the same domain share similar codon bias signatures (59), the use of our proposed regression models to predict the protein abundance of other species should be feasible. The extensive range of codon usage metrics used to train the machine learning algorithms could capture most patterns that underlie protein abundance in a domain. Since the models were trained using data from *S. cerevisiae*, this approach should work reasonably well for other eukaryotes. It might be possible to create similar predictive models for bacteria using absolute protein abundance data from *Escherichia coli* (69).

Our results underscore that codon usage metrics allow the prediction of protein abundances by machine learning. The observed difference between the CDSs of HAPs and the CDSs of LAPs supports this statement, as both groups sharply contrasted in the performed tests. Considering that codon usage bias is an intrinsic feature of gene sequences, all the metrics employed for compiling the data sets can be determined *in silico*, which simplifies the use of the proposed models for other species and is an advantage over previous attempts, which rely on other experimentally measured data. The machine learning models generated in our study can be a valuable tool for predicting protein abundances for yeasts that do not have large-scale quantitative proteomics available. Taking into account that the integration of protein abundances in GEMs allows improvement in phenotype simulations, our proposed regression models can be useful in system metabolic engineering approaches.

## DATA AVAILABILITY

All data and code are available in the GitHub repository: (https://github.com/LabFisUFV/protein_abundance_prediction)

Protein abundance data was obtained from Ho et al. (27) supplementary material: (https://www.cell.com/cell-systems/fulltext/S2405-4712(17)30546-X?#supplementaryMaterial)

Quantitative proteomics data for ecYeast7 simulations was obtained from Lahtvee et al. (57) supplementary material:

(https://www.cell.com/cell-systems/fulltext/S2405-4712(17)30088-1#supplementaryMaterial)

## SUPPLEMENTARY DATA

Supplementary Data are available online.

## ACKNOWLEDGEMENT

We thank Professor Leonardo Lopes Bhering (Department of General Biology, UFV) for granting access to the server at his laboratory (Laboratório de Processamento Biométrico (PROBIO)/BIODATA). We are grateful to Professor Marcelo Mendes Brandão (Institute of Biology, UNICAMP) and Dr. Otávio José Bernardes Brustolini (Laboratório Nacional de Computação Científica, Petrópolis) for their critical discussion and comments on this work.

