## Supplementary material for "Protein Abundance Prediction Through Machine Learning Methods"

**SUPPLEMENTARY INFORMATION**

**Expanded description of selected machine learning models**

We discovered that the AdaBoost estimator from Scikit-Learn, which was implemented with the TPOT-predicted stacked ensemble as a base estimator, was the best predictive model for the data set of all protein abundances and the data set of minimal medium abundances. The adaptive boosting algorithm (AdaBoost) employs a combination of regressors referred to as “weak learners” to generate a larger regressor, which is better than any single regressor. This combination of weak learners works by repeatedly training with different distributions of the input data set and combining their outputs. In the case of regression, the outputs are combined by a weighted average of median (1). While the weak learners are traditionally decision trees, the Scikit-Learn library supports the integration of other machine learning algorithms as weak learners to improve its performance, with the parameter referred to as the “base estimator”. The TPOT run exported a stacked ensemble of several algorithms (refer to Table S5). Thus, we decided to integrate the predicted pipeline into AdaBoost. We noticed that it outperformed all other algorithms and achieved higher $R^2$ scores and lower MAD values.

For the YPD medium data set, the best predictive model was the extreme gradient boosting algorithm from the XGBoost library, which was predicted and optimized by the H2O automated tool. XGBoost integrates multiple trees into a stronger learner, such as AdaBoost and other boosting algorithms. However, XGBoost has better performance than other algorithms as it is capable of running in parallel; does not need transformation of numerical, continuous data; and minimizes overfitting by implementing regularization procedures (2).

**SUPPLEMENTARY TABLES**

Table S1: List of codon usage metrics employed as features for constructing the training data sets.

| **Codon usage metrics** | **Reference** |
|---|---|
| Information theory-based codon usage bias (iCUB) | (3) |
| tRNA adaptation index (tAI) | (4) |
| Codon adaptation index (CAI) | (5) |
| Codon bias index (CBI) | (6) |
| Frequency of optimal codons (Fop) | (7) |
| Effective number of codons (ENC) | (8) |
| ENC alternative implementation (ENC’) | (9) |
| G+C content of gene | (10) |
| G+C of 3$^{rd}$ codon position | (10) |
| Base composition at silent sites | (10) |

| | |
|---|---|
| Hydropathicity of protein | (10) |
| Aromaticity of protein | (10) |
| B measure of codon bias | (11) |
| E measure of expression | (12) |
| Maximum likelihood codon bias (MCB) | (13) |
| Measure independent of length and composition (MILC) | (14) |
| MILC-based expression level predictor (MELP) | (14) |
| Synonymous codon usage orderliness (SCUO) | (15) |
| Gene codon bias (GCB) | (16) |
| Evolutionary selection pressure on nucleotide biosynthetic cost (Sc) | (17, 18) |

| | |
|---|---|
| Evolutionary selection pressure on gene translation efficiency (St) | (17, 18) |
| Nucleotide composition | (19) |

Table S2: List of features compiled for the training data sets using codon usage metrics calculated individually for gene and nucleotide composition numbers.

| | | | | | | | |
|---|---|---|---|---|---|---|---|
| iCUB | tAI | T3s | C3s | A3s | G3s | CAI_CODONW | CBI |
| Fop_CODONW | Nc | GC3s | GC | L_sym | L_aa | Gravy | Aromo |
| CAI_EMBOSS | CAI_coRdon | MELP | E | GCB | Fop_coRdon | MILC_SELF | MILC_REF |
| B_SELF | B_REF | MCB_SELF | MCB_REF | ENC | ENC_prime_SELF | ENC_prime_REF | SCUO |
| Sc | St | A | C | T | G | %A | %C |
| %T | %G | %G+C | %G+A | %G+T | %A+T | %A+C | %C+T |
| A1 | C1 | T1 | G1 | %A1 | %C1 | %T1 | %G1 |
| %G1+C1 | %G1+A1 | %G1+T1 | %A1+T1 | %A1+C1 | %C1+T1 | A2 | C2 |
| T2 | G2 | %A2 | %C2 | %T2 | %G2 | %G2+C2 | %G2+A2 |
| %G2+T2 | %A2+T2 | %A2+C2 | %C2+T2 | A3 | C3 | T3 | G3 |
| %A3 | %C3 | %T3 | %G3 | %G3+C3 | %G3+A3 | %G3+T3 | %A3+T3 |
| %A3+C3 | %C3+T3 | %G3s+C3s | | | | | |

Table S3: Example of how data sets were structured. The first column is the systematic name of all open reading frames (ORFs). The second column contains the protein abundance values for each ORF. From the third column to the last column, all columns are codon usage metrics or nucleotide composition numbers. A total of 91 columns are present in the data sets. The protein abundance values consist of the median values from several different quantitative proteomics analyses and are expressed as the number of molecules per cell.

| ORF | Protein abundance | iCUB | tAI | CAI (EMBOSS) | Fop | Sc | St | ... | %G3s+ C3s |
|---|---|---|---|---|---|---|---|---|---|
| Q0045 | 2440 | 20 | 0.263 | 0.584 | 0.599 | 0.05 | 0.04 | ... | 11.3 |
| Q0050 | 353 | 20 | 0.227 | 0.561 | 0.683 | 0.03 | -0.01 | ... | 9.9 |
| Q0055 | 271 | 20 | 0.242 | 0.601 | 0.679 | -0.01 | 0.02 | ... | 13.3 |
| Q0060 | 1029 | 20 | 0.199 | 0.587 | 0.810 | -0.17 | -0.01 | ... | 3.7 |
| Q0065 | 127 | 20 | 0.227 | 0.573 | 0.771 | -0.02 | 0.02 | ... | 6.2 |
| Q0085 | 1076 | 20 | 0.249 | 0.633 | 0.766 | 0.02 | -0.01 | ... | 7.3 |
| Q0115 | 183 | 25 | 0.196 | 0.600 | 0.805 | -0.01 | 0.06 | ... | 3.8 |
| ... | ... | ... | ... | ... | ... | ... | ... | ... | ... |
| YPR204W | 370 | 37 | 0.286 | 0.435 | 0.267 | 0.05 | 0.02 | ... | 47.9 |

Table S4: Regression evaluation metrics for all tested algorithms for each data set. Each algorithm was trained and evaluated by hold-out validation (75% training, 25% validation) using an independent data set for testing. Spearman's ρ and its associated p-value assesses the correlation between the predicted values and median values obtained by Ho et al. (20).

| **Algorithm** | **Coefficient of determination ($R^2$)** | **Median absolute deviation (MAD)** | **Spearman's ρ** | **p-value** |
|---|---|---|---|---|
| **All abundances data-set** | | | | |
| AdaBoost (Scikit-Learn) | 0.951 | 0.006 | 0.756 | 4.71E-101 |
| Random Forest (H2O) | 0.899 | 0.175 | 0.950 | 0 |
| Random Forest (Scikit-Learn) | 0.843 | 0.219 | 0.779 | 1.03E-220 |
| Bagging Meta-estimator (Scikit-Learn) | 0.842 | 0.188 | 0.750 | 9.11E-186 |
| Multilayer perceptron (H2O) | 0.834 | 0.224 | 0.916 | 0 |
| Extremely Randomized Trees (Scikit-Learn) | 0.775 | 0.277 | 0.778 | 2.62E-219 |
| Gradient Boosting Estimator (H2O) | 0.765 | 0.328 | 0.884 | 0 |
| Gradient Tree Boosting (Scikit-Learn) | 0.702 | 0.404 | 0.748 | 5.10E-184 |
| Ridge Regression (Scikit-Learn) | 0.643 | 0.435 | 0.769 | 1.64E-211 |
| Linear Regression (Scikit-Learn) | 0.642 | 0.441 | 0.770 | 1.53E-212 |
| Bayesian Ridge (Scikit-Learn) | 0.637 | 0.435 | 0.767 | 1.43E-209 |
| Theil-Sen (Scikit-Learn) | 0.633 | 0.455 | 0.765 | 1.43E-207 |
| Orthogonal Matching Pursuit (Scikit-Learn) | 0.631 | 0.439 | 0.753 | 4.45E-198 |
| Elastic Net | 0.626 | 0.456 | 0.753 | 1.54E-197 |

| | | | | |
|---|---|---|---|---|
| (Scikit-Learn) | | | | |
| Generalized Linear Model (H2O) | 0.625 | 0.453 | 0.801 | 9.36E-212 |
| Huber (Scikit-Learn) | 0.611 | 0.467 | 0.751 | 4.01E-196 |
| Lasso (Scikit-Learn) | 0.555 | 0.503 | 0.464 | 1.12E-58 |
| Nearest Neighbors (Scikit-Learn) | 0.487 | 0.544 | 0.521 | 5.77E-76 |
| Support Vector Regressor (Scikit-Learn) | 0.414 | 0.100 | 0.298 | 1.51E-23 |
| Lasso Lars (Scikit-Learn) | 0.020 | 0.815 | 0.732 | 6.92E-182 |
| Passive Agressive (Scikit-Learn) | -5.474 | 2.318 | 0.188 | 4.58E-10 |
| Gaussian Process (Scikit-Learn) | -11.518 | 1.389 | 0.114 | 4.39E-05 |
| XGBoost | -33.300 | 7.530 | 0.703 | 2.24E-161 |
| Decision Tree (Scikit-Learn) | -36.518 | 7.880 | 0.595 | 6.80E-99 |
| **Minimal medium data-set** | | | | |
| **AdaBoost (Scikit-Learn)** | **0.801** | **0.183** | **0.763** | **2.16E-100** |
| Random Forest (H2O) | 0.782 | 0.234 | 0.905 | 0 |
| Extremely Randomized Trees (Scikit-Learn) | 0.774 | 0.277 | 0.777 | 2.62E-219 |
| Bagging Meta-estimator (Scikit-Learn) | 0.772 | 0.232 | 0.749 | 6.44E-188 |
| Gradient Tree Boosting (Scikit-Learn) | 0.722 | 0.370 | 0.775 | 1.61E-217 |
| Gradient Boosting Estimator (H2O) | 0.716 | 0.329 | 0.856 | 8.69E-272 |

| | | | | |
|---|---|---|---|---|
| Multilayer perceptron (H2O) | 0.678 | 0.370 | 0.829 | 7.20E-240 |
| Random Forest (Scikit-Learn) | 0.645 | 0.386 | 0.750 | 1.16E-195 |
| Ridge Regression (Scikit-Learn) | 0.643 | 0.435 | 0.769 | 1.64E-211 |
| Linear Regression (Scikit-Learn) | 0.642 | 0.441 | 0.770 | 1.53E-212 |
| Bayesian Ridge (Scikit-Learn) | 0.637 | 0.435 | 0.767 | 1.43E-209 |
| Theil-Sen (Scikit-Learn) | 0.633 | 0.455 | 0.765 | 1.43E-207 |
| Orthogonal Matching Pursuit (Scikit-Learn) | 0.631 | 0.439 | 0.753 | 4.45E-198 |
| Elastic Net (Scikit-Learn) | 0.626 | 0.456 | 0.753 | 1.54E-197 |
| Generalized Linear Model (H2O) | 0.625 | 0.412 | 0.803 | 2.84E-213 |
| Huber (Scikit-Learn) | 0.611 | 0.467 | 0.751 | 4.01E-196 |
| Lasso (Scikit-Learn) | 0.555 | 0.503 | 0.732 | 6.92E-182 |
| Nearest Neighbors (Scikit-Learn) | 0.487 | 0.544 | 0.521 | 5.77E-76 |
| Passive Aggressive (Scikit-Learn) | 0.384 | 0.731 | 0.710 | 4.63E-166 |
| Lasso Lars (Scikit-Learn) | 0.020 | 0.815 | 0.464 | 1.12E-58 |
| XGBoost | -0.078 | 4530.561 | 0.778 | 1.12E-219 |
| Gaussian Process (Scikit-Learn) | -9.227 | 0.000 | 0.082 | 0.002612580833657 |
| Decision Tree (Scikit-Learn) | -37.463 | 8.007 | 0.616 | 2.32E-109 |
| **YPD medium data-set** | | | | |
| Extremely Randomized | 0.843 | 0.227 | 0.740 | 4.27E-178 |

| | | | | |
|---|---|---|---|---|
| Trees (Scikit-Learn) | | | | |
| Random Forest (H2O) | 0.828 | 0.265 | 0.917 | 0 |
| AdaBoost (Scikit-Learn) | 0.825 | 0.290 | 0.744 | 4.34E-181 |
| Random Forest (Scikit-Learn) | 0.816 | 0.282 | 0.743 | 2.06E-180 |
| Gradient Boosting Estimator (H2O) | 0.734 | 0.380 | 0.863 | 9.67E-282 |
| Multilayer perceptron (H2O) | 0.699 | 0.341 | 0.872 | 8.13E-294 |
| Gradient Tree Boosting (Scikit-Learn) | 0.699 | 0.407 | 0.748 | 3.55E-184 |
| Ridge Regression (Scikit-Learn) | 0.635 | 0.435 | 0.738 | 1.12E-176 |
| Orthogonal Matching Pursuit (Scikit-Learn) | 0.631 | 0.442 | 0.736 | 6.38E-175 |
| Linear Regression (Scikit-Learn) | 0.631 | 0.437 | 0.736 | 6.76E-175 |
| Elastic Net (Scikit-Learn) | 0.627 | 0.460 | 0.730 | 9.90E-171 |
| Bayesian Ridge (Scikit-Learn) | 0.626 | 0.456 | 0.734 | 6.65E-174 |
| Generalized Linear Model (H2O) | 0.624 | 0.465 | 0.802 | 1.01E-212 |
| Huber (Scikit-Learn) | 0.611 | 0.453 | 0.718 | 7.31E-163 |
| Lasso (Scikit-Learn) | 0.571 | 0.517 | 0.708 | 2.40E-156 |
| Decision Tree (Scikit-Learn) | 0.564 | 0.352 | 0.620 | 1.12E-109 |
| Nearest Neighbors (Scikit-Learn) | 0.490 | 0.553 | 0.499 | 1.19E-65 |
| Support Vector Regressor (Scikit-Learn) | 0.399 | 0.367 | 0.286 | 1.18E-20 |

| | | | | |
|---|---|---|---|---|
| Passive Aggressive (Scikit-Learn) | -0.723 | 0.904 | 0.389 | 2.15E-38 |
| Bagging Meta-estimator (Scikit-Learn) | -0.795 | 1.834 | 0.756 | 4.39E-190 |
| Gaussian Process (Scikit-Learn) | -11.518 | 1.389 | 0.114 | 4.39E-05 |
| XGBoost (Scikit-Learn) | -33.301 | 7.532 | 0.690 | 1.88E-145 |

Table S5: List of algorithms that are part of the stacked ensemble applied as weak learners for the AdaBoost estimator. This list was generated as part of the TPOT automated pipeline prediction. The algorithms are utilized in this order.

| **Algorithm** | **Scikit-Learn/XGBoost function** |
|---|---|
| Cross-validated Lasso | LassoLarsCV |
| Linear Support Vector Regression | LinearSVR |
| Ridge regression with built-in cross-validation | RidgeCV |
| Linear Support Vector Regression | LinearSVR |
| Extremely Randomized Trees Regressor | ExtraTreesRegressor |
| Ridge regression with built-in cross-validation | RidgeCV |
| Linear Support Vector Regression | LinearSVR |
| Ridge regression with built-in cross-validation | RidgeCV |
| Stochastic Gradient Descent | SGDRegressor |
| Linear Support Vector Regression | LinearSVR |
| Elastic Net model with iterative fitting along a regularization path | ElasticNetCV |
| Extreme Gradient Boosting | XGBRegressor |
| Extremely Randomized Trees Regressor | ExtraTreesRegressor |
| Linear Support Vector Regression | LinearSVR |
| Random Forest Regressor | RandomForestRegressor |

**SUPPLEMENTARY FIGURES**

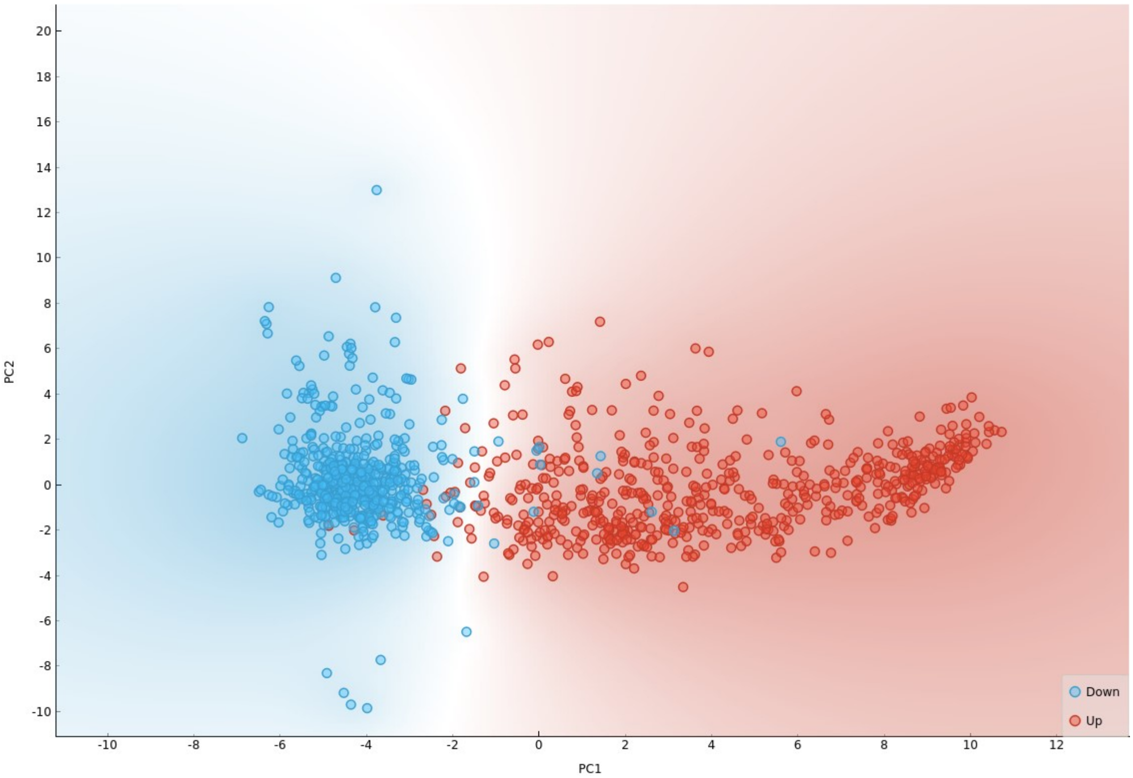


Figure S1: Principal component analysis of RSCU values calculated from CDSs of highly abundant proteins (HAP) and lowly abundant proteins (LAP). Two distinct groups of CDSs could be observed. The first group is composed of mostly HAP, and the second group is composed of mostly LAP.

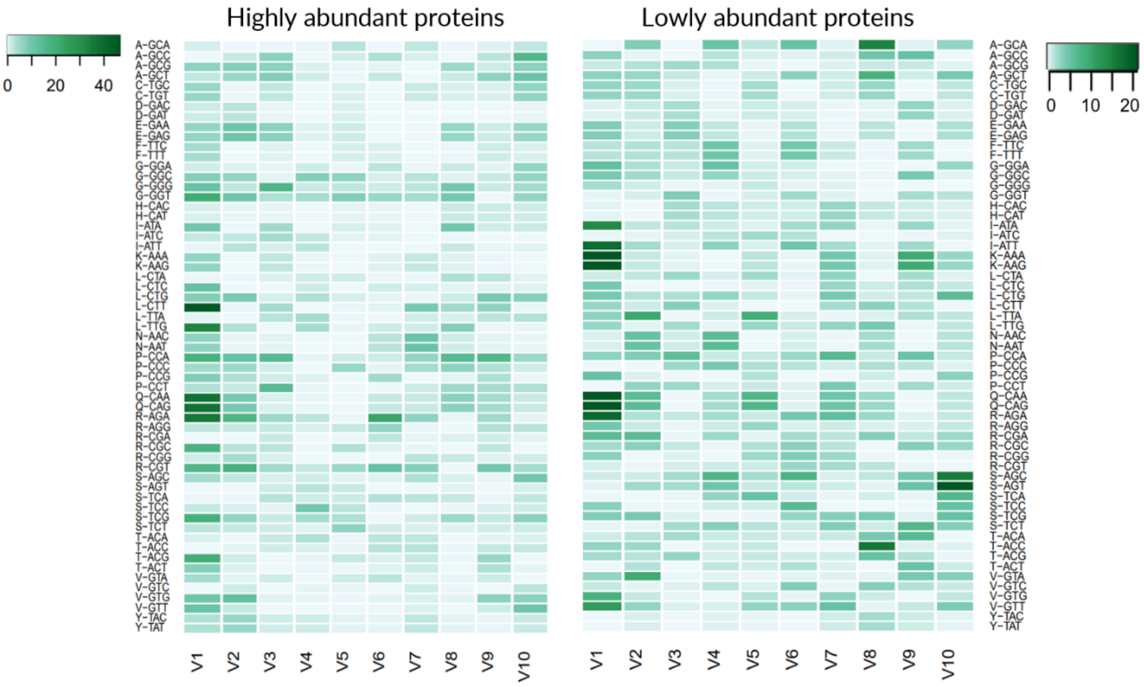


Figure S2: Evolutionary selection for position-dependent codon usage bias as determined by the chi-squared test on the matrix constructed with the CodG package. Each CDS was equally divided into 10 bins to evaluate how each position contributes to the overall codon usage. A) Deviation from uniformity in the CDSs of highly abundant proteins show bias towards 5’ end. B) The CDSs of lowly abundant proteins show higher uniformity.

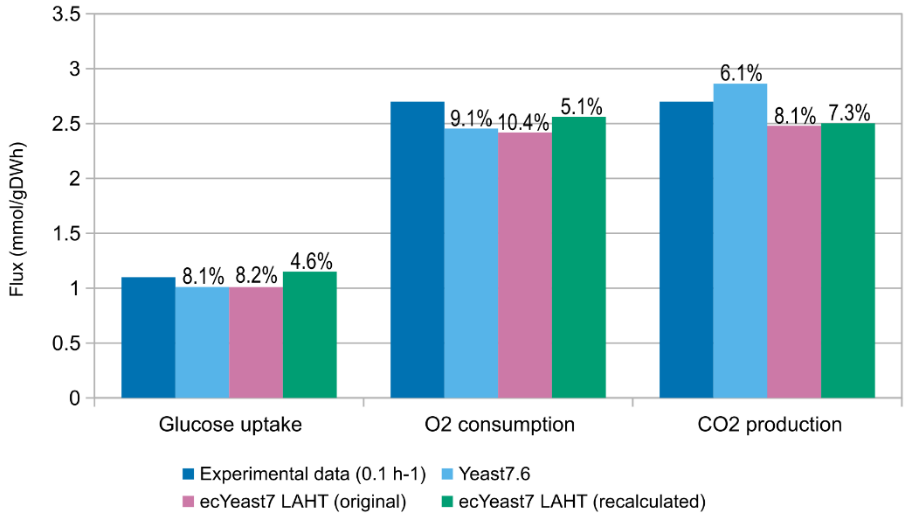


Figure S3: Predictions of metabolic flux obtained by the Yeast7 model and ecYeast7. We attempted to validate our unit conversion step by replicating the analysis performed by Sánchez et al. (2017). The "original" ecYeast7 LAHT model employs the quantitative proteomics data from Lahtvee et al. (2016). The "recalculated" ecYeast7 LAHT model applies the median absolute values from Ho et al. (2018). Percentage values represent the relative error when compared to experimental values
